# Early life pain experience changes adult functional pain connectivity in the rat somatosensory and the medial prefrontal cortex

**DOI:** 10.1101/2022.03.01.482554

**Authors:** Pishan Chang, Lorenzo Fabrizi, Maria Fitzgerald

## Abstract

Early life pain experience (ELP) alters adult pain behaviour and increases injury induced pain hypersensitivity, but the effect of ELP upon adult functional brain connectivity is not known. We have performed continuous local field potential (LFP) recording in the awake adult male rats to test the effect of ELP upon functional cortical connectivity related to pain behaviour. Somatosensory cortex (S1) and medial prefrontal cortex (mPFC) LFPs evoked by mechanical hindpaw stimulation were recorded simultaneously with pain reflex behaviour for 10 days after adult skin injury. We show that, post adult injury, S1 LFP delta and gamma energy and S1 LFP delta/gamma frequency modulation are significantly increased in ELP rats compared to controls. Adult injury also induces increases in S1-mPFC functional connectivity which is significantly prolonged in ELP rats, lasting 4 days compared to 1 day in controls. Importantly, the increases in LFP energy and connectivity in ELP rats were directly correlated with increased behavioural pain hypersensitivity. Thus, early life pain (ELP) alters adult brain functional connectivity, both within and between cortical areas involved in sensory and affective dimensions of pain. The results reveal altered brain connectivity as a mechanism underlying the effects of early life pain upon adult pain perception.

**Significance Statement:** Pain and stress in early life has a lasting impact upon pain behaviour and may increase vulnerability to chronic pain in adults. Here we record pain-related cortical activity and simultaneous pain behaviour in awake adult male rats previously exposed to pain in early life. We show that functional connectivity within and between the somatosensory cortex and the medial prefrontal cortex is increased in these rats and that these increases are correlated with their behavioural pain hypersensitivity. The results reveal that early life pain alters adult brain connectivity, which may explain the impact of childhood pain upon adult chronic pain vulnerability.

## Introduction

Exposure to pain and injury in early life pain (ELP) is associated with altered pain behaviour in adults. Evidence from both human and animal studies shows that repeated painful procedures or surgical incision during a critical period of early postnatal development has significant long-term effects on pain processing (Walker et al., 2009b, 2009a; Beggs et al., 2012; Schwaller and Fitzgerald, 2014; van den Hoogen et al., 2018). The mechanisms underlying the effects of ELP involve changes in spinal cord nociceptive circuitry (Torsney and Fitzgerald, 2003; Li and Baccei, 2019), early life spinal microglial activation (Moriarty et al., 2019) and altered descending brain stem pain control (Walker et al., 2015). There is also evidence from human studies of structural changes in the thalamus and cortex (Duerden et al., 2018) and functional changes in descending pain control from supraspinal sites (Walker et al., 2018). The importance of this extends into a wider area of the long-term consequences of early life stress and pain which, by inducing long-term alterations in brain function and behaviour may lead to higher susceptibility to chronic pain (Jones et al., 2009; Denk et al., 2014; Melchior et al., 2021; Ririe et al., 2021). However, as yet, there is no evidence that ELP has any effect upon adult cortical pain networks or upon functional connectivity between the key cortical regions involved in the sensory and emotional dimensions of pain.

A wide network of brain areas is involved in acute pain processing, including primary (S1) and secondary (S2) somatosensory cortices, the medial prefrontal cortex (mPFC), insula, thalamus, and prefrontal areas (Apkarian et al., 2005; Duerden and Albanese, 2013; Tan and Kuner, 2021). To address whether ELP impacts upon cortical function from the sensory-discriminative and emotional/cognitive perspectives, the S1 and mPFC are attractive targets (Tan and Kuner, 2021). S1 is a functionally defined part of the somatosensory and nociceptive system and processes sensory nociceptive information about pain from an early age in both rodents and humans (Chang et al., 2016, 2020b; Jones et al., 2021). S1 encodes nociceptive intensity and perceived pain intensity (Mancini et al., 2012) and gamma-band oscillations in this area correlate with subjective pain perception (Ong et al., 2019). While mPFC is critically involved in numerous cognitive functions (Euston et al., 2012; Chang et al., 2020a) and emotion behaviour (Cao et al., 2018; Huang et al., 2020), this area also plays an important role in the emotional and affective aspects of pain, and could modulate pain sensation by controlling the flow of afferent sensory stimuli into the dorsal horn through descending control pathways (Zhang et al., 2015; Huang et al., 2020). Here we hypothesise that ELP alters pain related connectivity in the adult S1 and mPFC and that this is associated with increases in adult pain related behaviour.

In this study, we used a well-established model of injury and postoperative pain: hind-paw plantar incision of skin and underlying muscle (Brennan et al., 1996; Beggs et al., 2012) to examine the impact of ELP upon pain behaviour and associated neural activity in S1 and mPFC. We recorded local field potentials (LFPs) in S1 and mPFC in awake, freely moving adult rats and analysed the oscillatory energy within those LFPs and the functional connectivity within and between these areas. Acute pain is associated with defined changes in cortical oscillations (Tan et al., 2021). In humans, gamma-band oscillations in S1 correlate with subjective pain perception (Heid et al., 2020; Yue et al., 2020) and are strengthened in rodent S1 cortex during nociception and inflammatory pain in association with behavioural nociceptive hypersensitivity (Tan et al., 2019). We also analyse phase-amplitude coupling and coherence of neuronal oscillations as putative mechanisms of regional and inter-areal communication (Buzsaki, 2004; Peng and Tang, 2016). Together, our results provide new insights into how early life pain (ELP) alters adult cortical function underlying sensory and emotional dimensions of pain behaviour.

## Materials and Methods

### Experimental animals

All experiments were performed in accordance with the United Kingdom Animal (Scientific Procedures) Act 1986. Reporting is based on the ARRIVE Guidelines for Reporting Animal Research developed by the National Centre for Replacement, Refinement and Reduction of Animals in Research, London, United Kingdom. Male Sprague-Dawley rat pups were obtained from the Biological Services Unit, University College London. Rats were housed in cages of five age-matched animals (>P21) or with the dam and littermates (P) 3 to 21 under controlled environmental conditions (24–25°C; 50–60% humidity; 12 h light/dark cycle) with free access to food and water. In the case of rat pups, handling and maternal separation were kept to a minimum. All animals were exposed to the same standard caging, handling and diet throughout development. The different experimental groups are represented in Figure 1A and protocol for probing the impact of nociceptive inputs in the early life on central pain processing and adult pain sensitivity is summarized in Figure 1B

**Figure 1.**
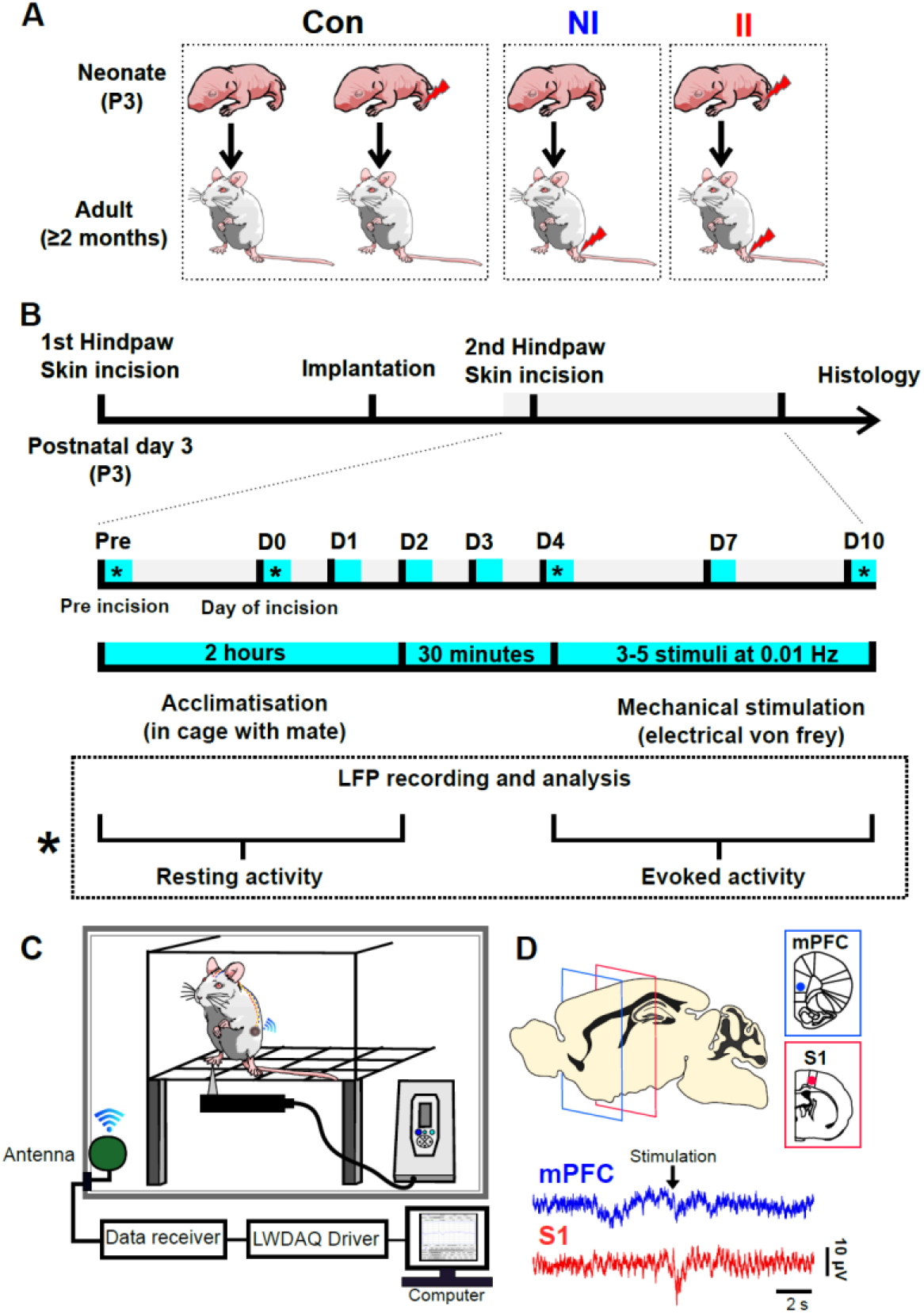
Experiment design. (A) Schematic of experimental groups. II: neonatal incision on postnatal day 3 and repeat incision 2 months later in adulthood (ELP model). NI: littermate control with equivalent anaesthesia, handling and maternal separation on post-natal day 3 and first incision in adulthood. Con: pooled data from animals having neonatal incision only and from age-matched non-incised litter mates from the same colony. (B) Experimental protocol for probing the impact of early life pain (ELP) on adult cortical pain processing and pain behaviour. Upper scale: timeline for recording cortical local field potentials (LFP) and pain behaviour, where * marks days of simultaneous electronic von Frey hair (eVF) stimulation and LFP recording. Lower box: Detail of testing protocol for recording resting LFPs and eVH evoked LFP recording. (C) Schematic of the experimental set-up for simultaneous recording of neural LFP activity in medial prefrontal cortex (mPFC) and primary somatosensory cortex (S1) in awake adult rats using wireless telemetry while applying electronic von Frey hairs (eVF) to the plantar hindpaw. (D) Sample traces of simultaneous S1 and mPFC evoked potentials evoked by mechanical eVF stimulation of the plantar hindpaw.

### Plantar hind-paw incision

Male rat pups on postnatal Day 3 were anaesthetized and plantar hindpaw incision performed. Under general anesthesia with 2% isoflurane in 100% oxygen (flow rate, 1–1.5 L/min), a midline longitudinal incision was made through the skin and fascia extending from the midpoint of the heel to the proximal border of the first footpad and the underlying plantar muscle elevated and incised. The same relative length of incision was performed in adult animals as previously described (Brennan et al., 1996; Walker et al., 2009b). Skin edges were closed with 5–0 nylon suture (Ethicon). The whole procedure took 3–5 min. After plantar hindpaw incision, rats were placed in a recovery chamber and allowed to recover from the general anesthesia before returning to their home cage.

Four experimental groups were used: **II:** neonatal incision on postnatal day 3 and repeat incision 2 months later in adulthood. **NI:** littermate control with equivalent anaesthesia, handling and maternal separation on post-natal day 3 and having incision in adulthood. Animals having neonatal incision and follow-up in adulthood **(IN)** and age-matched non-incised litter mates from the same colony **(NN)** were pooled data and used as control group **(Con)**, due to that there is no significant different between the two groups (Figure 1A)

### Pain hypersensitivity testing

To test behavioural pain hypersensitivity following hind-paw skin incision, an electronic von Frey unit (EVF4, Bioseb) was used to measure hindpaw mechanical flexion withdrawal thresholds (Ferrier et al., 2015, 2016). Following habituation for 30 min on an elevated mesh platform, a mechanical stimulus was applied to the plantar surface of the hindpaw adjacent to the distal half of the incision. The von Frey (eVF) apparatus, which has a measurement range of 0-500 g with 0.1 g resolution, consists of a plastic tip fitted in a hand-held force transducer, which was applied to the rat hindpaw from below with force (g) gradually increased until paw withdrawal. The force that induced paw withdrawal was digitized and recorded automatically by the unit and used as the threshold for mechanical nociception. For each recording session, the eVF was applied 3-5 times at ∼50 sec intervals. Simultaneous recording from both S1 and mPFC accompanied testing of eVF withdrawal thresholds (Figure 1 C, D).

### Surgical Preparation and Transmitter Implantation for Long-term Recording

Rat were anaesthetised with 2.5-3 % isoflurane (Abbot, AbbVie Ltd., Maidenhead, UK) in 100% oxygen (flow rate of 1-1.5 litre/min) via gas anaesthesia mask (Model 906, David Kopf Instruments) from a recently calibrated vaporizer (Harvard Apparatus, Cambridge, MA). Body temperature was maintained with a heat blanket during surgery. A transmitter (A3028D-DDA, Open Source Instruments, Brandeis, Boston, USA)(Chang et al., 2011) was implanted subcutaneously with the depth recording electrodes (J-electrode (wire 125-μm dia 316SS 10kOhm impedance), a Teflon-insulated stainless steel electrode, Open Source Instruments, Brandeis, Boston, USA) positioned in mPFC (3.2 mm anterior, 0.5 mm lateral, 4 mm ventral) and primary somatosensory hindpaw cortex (1 mm posterior, 2.5 mm lateral, 2 mm ventral) (Paxinos et al., 2013; Chang et al., 2016). The reference electrode was implanted over the cerebellum posterior to lambda. The whole assembly was held in place with dental cement (Simplex Rapid, Acrylic Denture Polymer, UK). A subcutaneous injection of bupivacaine and metacam was provided for post-surgical pain management. At the end of surgery, enrofloxacin (5mg/kg, Baytril, Bayer health care) and pre-warm saline (0.5-1 ml) were administered subcutaneously. The animals were placed in a temperature controlled (25°C) recovery chamber until ambulatory and closely monitored at least 1-2 hours before returning to their home cage to allow recovery for at least 14 days after surgery.

The transmitter, which has no adverse effects (Chang et al., 2016), was implanted for data recordings. During all recording sessions, continuous LFP recordings were recorded (bandpass filter: 0.2 Hz to 160 Hz, 512Hz sampling rate with 16 bit resolution) using LWDAQ Software (Open Source Instruments, Brandeis, Boston, USA). Animals were carefully monitored daily and were euthanized at the end of experiment with carbon dioxide (CO_2_). The brain was removed and immediately immersed in 4% paraformaldehyde for >24 hours before being transferred to 30% sucrose post-fixation solution. Brain sections (40-μm thick thickness) were cut using a microtome (Leica SM2000R, Leica Microsystems (UK) Ltd., United Kingdom) and stained with Cresyl violet to allow histological location of the electrode track. This procedure allowed us to verify recording electrode locations, and LFP data were only included in the study if electrode tips were located in mPFC and S1 (Figure 1D).

### Analysis of electrophysiology data

Data analysis was performed with Brainstorm (Tadel et al., 2011), which is free and open source for electrophysiology data visualization and processing through a simple and intuitive graphical user interface (GUI) (http://neuroimage.usc.edu/brainstorm) and custom Matlab scripts (The Mathworks Inc.MA, USA).

### Evoked LFP data processing

#### LFP Pre-processing

For our initial analyses, continuous LFP recordings from each region were segmented into 10s epochs that lasted from 5s before to 5s after the peak of evoked LFP. Each epoch was visually inspected for artefacts prior to further analysis. Any epochs that exhibited artefacts during visual inspection were excluded from subsequent analysis.

#### Time-frequency Analysis

Activity changes in LFP in different frequency bands were calculated using the Hilbert transform (Le Van Quyen et al., 2001; Bruns, 2004; Tadel et al., 2011). Each epoch was filtered in various frequency bands with bandpass filters for delta (2–4 Hz), theta (4–8 Hz), alpha (8– 12 Hz), beta (12–30 Hz), and gamma (30–90 Hz) band. The magnitude (µV/sqrt(Hz)) of the Hilbert transform of a narrow-band signal is a measure of the envelope of this signal, and therefore gives an indication of the activity in this frequency band. The energy magnitude data were then averaged across repetitions within each animal. Stimulus-induced changes in energy magnitude for each animal were then calculated by normalized to mean of baseline (−4 to -1s).

#### Time-resolved Phase-amplitude Coupling Analysis

This approach measures cross-frequency coupling between bursts of high-frequency oscillations and the phase of lower frequency rhythms, over a time-window, which slides along the electrophysiological data (Samiee and Baillet, 2017). mPFC and S1 time-courses were examined for changes in phase of slow oscillation at delta band (2-4 HZ) coupled to the amplitude of a faster rhythm at gamma (30-90 Hz) band. Phase and amplitude information were obtained via the Hilbert transform. The coupling between phase and amplitude was then quantified and Modulation Index values were calculated. To avoid edge artefacts, which can result in spurious phase-amplitude coupling (PAC) (Kramer et al., 2008), the first 2 s and last 2 s of each trial was used as buffer. These were then averaged across repetitions within each animal. Stimulus-induced Phase-Amplitude Coupling for each animal were then calculated by normalized to mean of baseline (−2.5 to -1 s)

#### Time-resolved Phase locking Analysis

To evaluate the functional connectivity between mPFC and S1, we estimated phase-locking value between the LFPs simultaneously recorded at the two areas in different frequency bands (Lachaux et al., 1999). To do this we (i) band-pass filtered the LFPs at S1 and mPFC in the delta (2–4 Hz), theta (4–8 Hz), alpha (8–12 Hz), beta (12–30 Hz), and gamma (30–90 Hz) frequency bands; (ii) applied Hilbert transform to the band-passed signals; (iii) calculate the instantaneous phase-locking value between mPFC and S1. PLVs were then averaged across repetitions within each animal. Stimulus-induced magnitude changes in LFP energy for each animal were then calculated by normalized to mean of baseline (−4 to -1s)

### Quantification and Statistical Analysis

Detailed statistical analysis was performed using GraphPad Prism 6 (GraphPad Software), SPSS (Statistical Product and Service Solutions, IBM). All data are presented as mean ± SEM. Comparisons of means were performed using one way ANOVA with Tukey post hoc test if the data were normally distributed; Kruskal-Wallis test with post hoc Dunn’s multiple comparisons test if the data were not normally distributed (with the Shapiro-Wilk test used to assess normality of the data distributions). Generalized linear model (GLM) Type III tests followed by Bonferroni post hoc tests were used for analysis of repeated-measures behaviour data. Differences were considered statistically significant at p < 0.05.

Estimation statistics (open source estimation program available on https://www.estimationstats.com)(Ho et al., 2019) were used to compute the change of electrophysiological data in mPFC/S1 in response to eVF stimulation with days following injury (D0, D4, and D10), compared to pre-injury activity. Mean differences are shown using Cumming estimation plots, with each graphed as a bootstrap sampling distribution (5000 bootstrap samples). The p value(s) reported are the likelihood(s) of observing the effect size(s), if the null hypothesis of zero difference is true. For each permutation p value, 5000 reshuffles of the control and test labels were performed; p < 0.05 is considered a significant difference.

Pearson correlation was applied to calculate the correlation between pain sensitivity and electrophysiological data. The significance threshold for all correlation tests was set at p < 0.05.

## Results

### Early life pain (ELP) increases injury induced hyperalgesia and pain in adult life

Behavioural pain threshold testing confirmed the impact of early life pain upon adult pain behaviour, as described previously (Beggs et al., 2012; Moriarty et al., 2019). We measured the amplitude and duration of hindlimb withdrawal reflexes in response to electronic von Frey hair stimulation following skin injury in adult male rats. Figure 1 shows von Frey hair pain thresholds in adult ELP male rats before and 10 days after an adult hindpaw skin incision **(II)**. This is compared to age-matched animals with no ELP, experiencing their first hindpaw skin incision in adulthood **(NI)** and control rats that have ELP only or no incisions at all **(Con)** (Figure 1).

Hindpaw skin injury caused von Frey hair thresholds to fall in both groups of adult rats (NI and II) compared to the control (Con) group, indicating a significant post injury pain and hyperalgesia. Consistent with previous reports (Beggs et al., 2012; Moriarty et al., 2019), animals that experienced early life pain (ELP) (II, n=9) developed significantly lower paw withdraw thresholds (PWT), compared to injured animals with no ELP (NI, n=10). In addition, as in earlier studies (Walker et al., 2009b), ELP resulted in more prolonged as well as enhanced hyperalgesia, lasting up to 10 days after hindpaw skin incision, compared to 3-4 days in non ELP rats.

### Early life pain (ELP) increases post injury evoked delta and gamma activity in adult S1

To test the impact of early life pain experience upon pain related neural activity in S1 and mPFC, we next investigated evoked potentials (EP) and oscillatory neural activity in S1 and mPFC evoked by von Frey hair stimulation following skin injury. Evoked potential (EP) amplitudes in S1 and mPFC did not differ between groups (enter data in here), and so to gain further insight into the pattern and time-course of evoked cortical activity following skin injury, the EP energy was analysed in delta (2–4 Hz), theta (4–8 Hz), alpha (8–12 Hz), beta (12–30 Hz), and gamma (30–90 Hz) frequency bands. Fig 3 & Fig 4 show a significant increase in the vFh evoked delta energy (Fig 3C,D) and gamma energy (Fig 4C,D) in S1 following skin incision in ELP rats, which is not observed in the other adult rat groups, NI and Con. These increases in delta and gamma energy in ELP rats were recorded in the 2-3 hours post injury (D0) and had recovered by 4 days post injury. They were not observed in mPFC.

**Figure 2:**
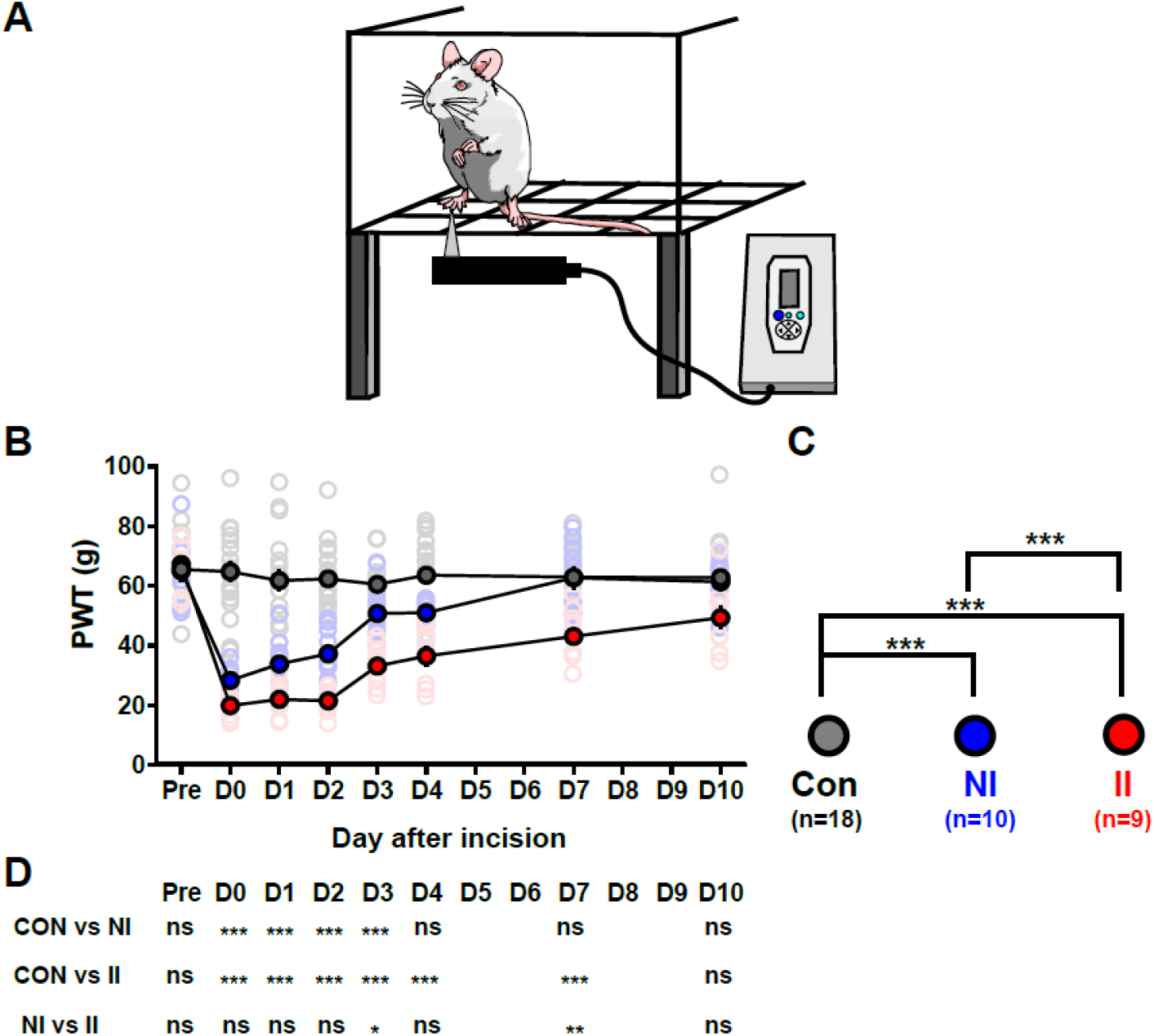
Early life pain (ELP) increases hyperalgesia following skin incision injury in adult rats. (A) Electronic von Frey hair (eVF) testing of the plantar skin adjacent to the wound (B) Plot of contralateral mechanical paw withdrawal thresholds (PWT), before (Pre) and up to 10 days (D) after hindpaw skin incision in adult rats. Mean ± SEM with individual data superimposed. (B) Statistical differences between groups using generalized linear models (C) Summary the post-hoc pairwise comparisons with Bonferroni correction. *P<0.05, **P < 0.01 ***p < 0.001. Non-incised controls (Con, n=18), incision in adults without neonatal incision (NI, n=10) and incision in adults with neonatal incision (II, n=9).

**Figure 3:**
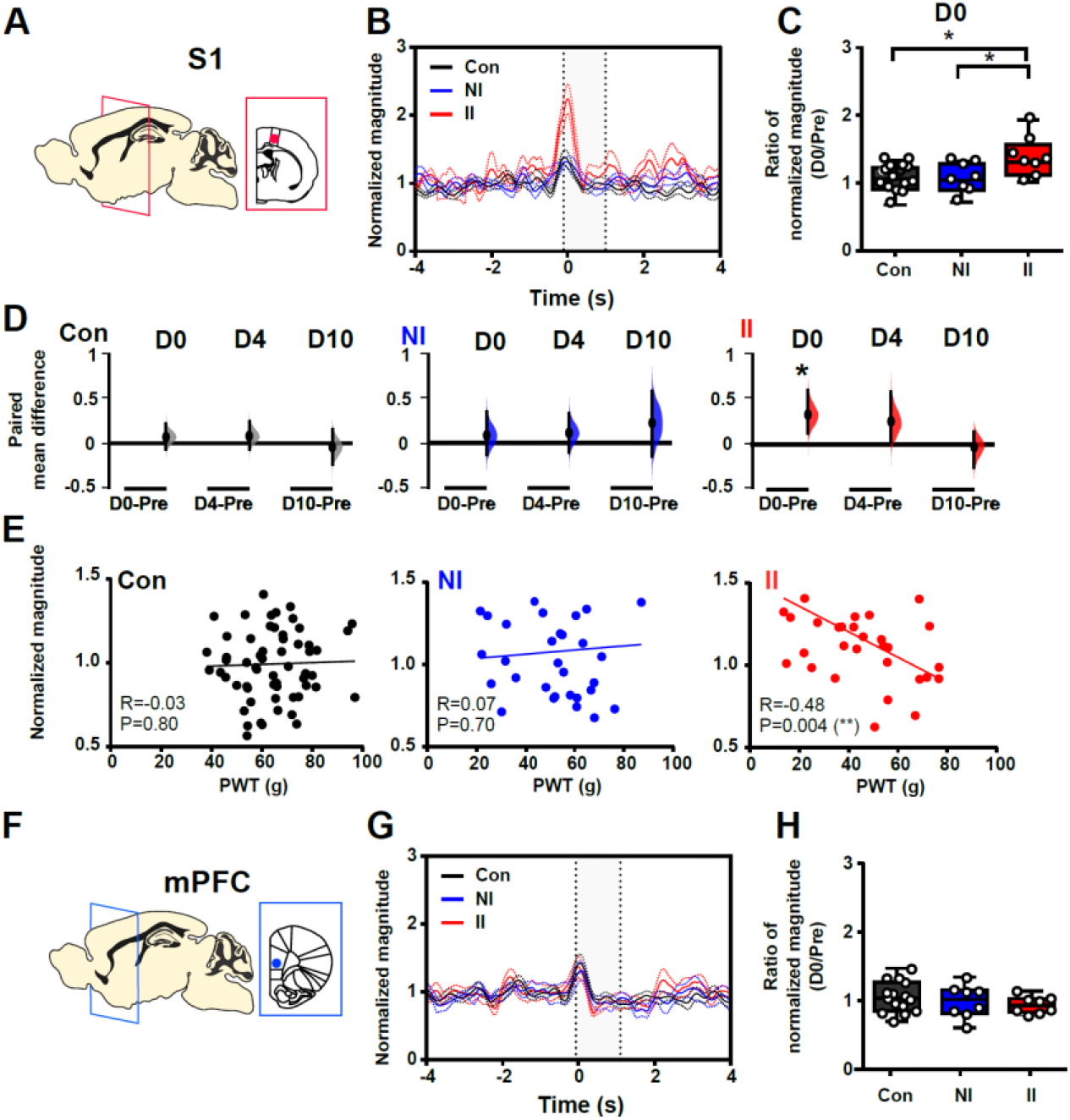
Early life pain (ELP) increases stimulus evoked delta energy in adult S1 post injury. Mechanical (eVF) stimulation of the hindpaw following injury in ELP adult rats (II, red), non-ELP rats (NI, blue) and controls (Con, black) while recording evoked potentials in the somatosensory cortex (S1) (B) Time-frequency (TF) magnitude (µV/sqrt(Hz)) in delta frequency (2-4 Hz) oscillations were normalized to mean of baseline (−4 to -1 s) and presented as normalized magnitude (mean ± SEM). (C) Comparing the pain-induced changes in evoked S1 delta activity expressed as normalized magnitude (D) evoked S1 delta energy following injury (D0), 4 days (D4) and 10 days (D10) after injury expressed ad paired mean difference with pre-injury (Pre). The paired mean difference for comparisons are shown as Cumming estimation. Each paired mean difference (D0-Pre: D0 minus Pre; D4-Pre: D4 minus Pre; D10-Pre: D10 minus pre) is plotted as a bootstrap sampling distribution; 95% confidence intervals are indicated by the ends of the vertical error bars. Statistical analysis was performed using permutation t-test (Randomization: 5000). (E) Correlations between paw-withdraw threshold (PWT) and evoked S1 delta activity (normalized magnitude). The scatter plots represent the correlations PWT and normalized TF magnitude (Pre to D10) with continuous line showing the linear regression. Pearson correlation coefficient (R) with significance (P value) is presented in the figures. Non-incised adult controls (Con, n=15), incision in adults without neonatal incision (NI, n=8) and incision in adults with neonatal incision (II, n=8).

**Figure 4:**
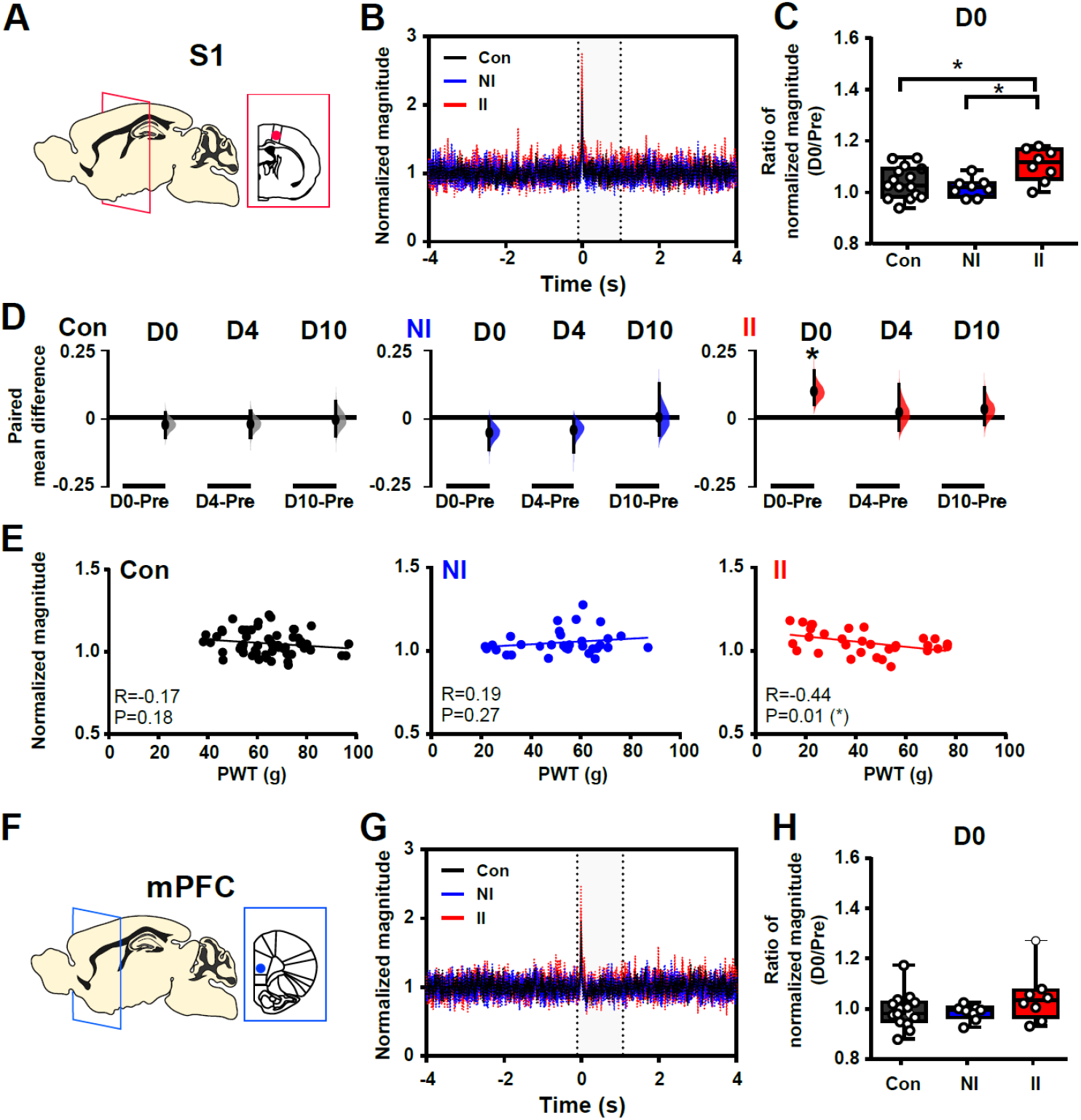
Early life pain (ELP) increases stimulus evoked gamma energy in adult S1 post injury. Mechanical (eVF) stimulation of the hindpaw following injury in ELP adult rats (II, red), non-ELP rats (NI, blue) and controls (Con, black) while recording evoked potentials in the somatosensory cortex (S1) (B) Time-frequency (TF) magnitude (µV/sqrt(Hz)) in gamma frequency (30-90 Hz) oscillations were normalized to mean of baseline (−4 to -1 s) and presented as normalized magnitude (mean ± SEM). (C) Comparing the pain-induced changes in evoked S1 gamma activity expressed as normalized magnitude (D) evoked S1 gamma energy following injury (D0), 4 days (D4) and 10 days (D10) after injury expressed ad paired mean difference with pre-injury (Pre). The paired mean difference for comparisons are shown as Cumming estimation. Each paired mean difference (D0-Pre: D0 minus Pre; D4-Pre: D4 minus Pre; D10-Pre: D10 minus pre) is plotted as a bootstrap sampling distribution; 95% confidence intervals are indicated by the ends of the vertical error bars. Statistical analysis was performed using permutation t-test (Randomization: 5000). (E) Correlations between paw-withdraw threshold (PWT) and evoked S1 delta activity (normalized magnitude). The scatter plots represent the correlations PWT and normalized TF magnitude (Pre to D10) with continuous line showing the linear regression. Pearson correlation coefficient (R) with significance (P value) is presented in the figures. Non-incised adult controls (Con, n=15), incision in adults without neonatal incision (NI, n=8) and incision in adults with neonatal incision (II, n=8).

Importantly, the magnitude of S1 evoked delta and gamma activity was significantly correlated to pain sensitivity, or fall in behavioural von Frey hair paw withdrawal threshold (PWT), as indicated by the inverse correlation of S1 delta power (Fig 3E) and S1 gamma power (Fig 4E) with PWT in II male adult rats.

### Early life pain (ELP) increases post injury evoked delta-gamma modulation in adult S1

Since delta and gamma energy evoked by mechanical stimulation (eVF) post injury is increased in S1 in ELP rats, we next asked whether ELP altered cross-frequency modulation (delta (2-4Hz) vs. gamma (30-90 Hz)) associated with the observed differences in pain sensitivity following hindpaw skin incision. Cross-frequency coupling potentially provides a mechanism for investigating local-to-wide networks synchronization and interaction (Florin and Baillet, 2015). Here, in order to evaluate event related changes in phase-amplitude coupling, we used time-resolved phase-amplitude coupling (tPAC). Fig 5 shows a significant enhancement of evoked phase amplitude modulation between delta and gamma in S1 evoked eVF potentials following hindpaw skin injury (D0) in II rats (Figure 5 A-C). This increase in delta-gamma coupling was not seen in mPFC (Figure 5D). The enhanced evoked delta-gamma modulation in S1 was observed on the day of injury and return to pre injury levels by 4 days (D4) after skin incision in II rats. There was no significant alteration in S1 evoked delta-gamma modulation in NI and Con rats (Figure 5E). To determine whether this increase in evoked S1 delta-gamma coupling is associated with the enhanced pain sensitivity, we subsequently examined the correlation between the two measures. A significant inverse correlation was found between delta-gamma coupling and PWT in II rats, but not in NI and Con rats (Figure 5F). Thus, pain related stimulus evoked delta-gamma coupling in the somatosensory cortex and its association with pain behaviours is selectively increased in adult ELP rats.

**Figure 5:**
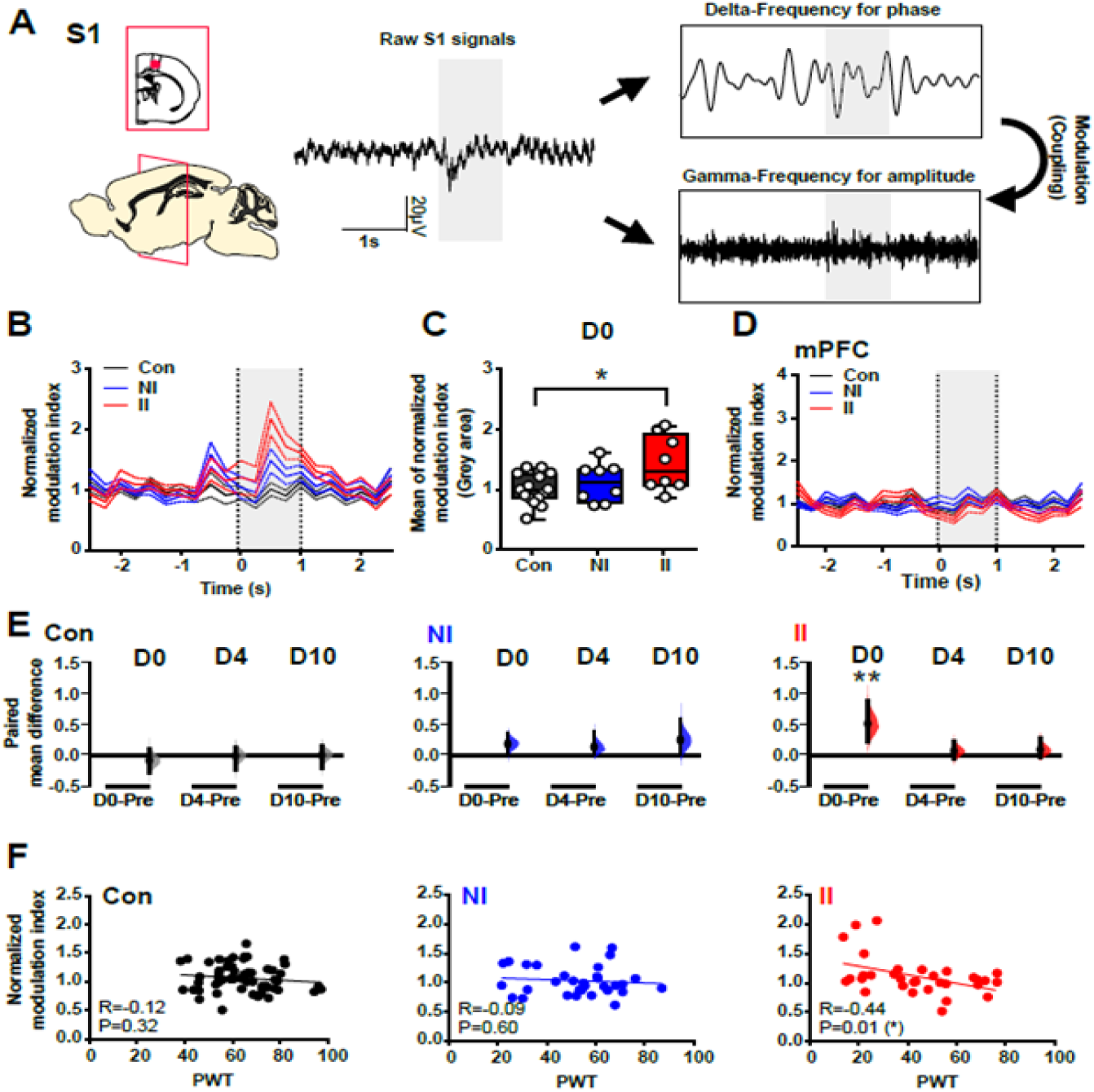
Early life pain (ELP) increases evoked cross-frequency modulation between delta (phase) and gamma (amplitude) in adult S1 post injury. Mechanical (eVF) stimulation of the hindpaw following injury in ELP adult rats (II, red), non-ELP rats (NI, blue) and controls (Con, black) while recording evoked potentials in the somatosensory cortex (S1). (A) Sample trace of potential in S1 evoked by hindpaw electronic von Frey hair (eVF) and diagram illustrating the principle of modulation coupling. (B) Time resolved phase-amplitude coupling values in S1 were normalized to mean of baseline (−2.5 to -1 s) and present as normalized modulation index (mean ± SEM). (C) The normalized modulation index during the evoked response (grey area, -0.02s to 1 s) were compared using one-way ANOVA with Tukey’s multiple comparisons test. Data are presented as box-whisker plots indicating the median, 25-75th percentiles, and minimum-maximum values with data for individual rat superimposed. (D) Time resolved phase-amplitude coupling values in mPFC show no increase. (E) Evoked S1 delta-gamma coupling following injury (D0), 4 days (D4) and 10 days (D10) after injury were compared to pre-injury (Pre). The paired mean difference for comparisons are shown as Cumming estimation. Each paired mean difference (D0-Pre: D0 minus Pre; D4-Pre: D4 minus Pre; D10-Pre: D10 minus pre) is plotted as a bootstrap sampling distribution; 95% confidence intervals are indicated by the ends of the vertical error bars. Statistical analysis was performed using permutation t-test (Randomization: 5000).(F) Correlations between paw-withdraw threshold (PWT) and delta-gamma modulation in S1 expressed as normalized modulation index. The scatter plots represent the correlations PWT and normalized modulation index with continuous line as linear regression. Pearson correlation coefficient (R) with significance (P value) *p < 0.05, **P<0.01. Non-incised adult controls (Con, n=15), incision in adults without neonatal incision (NI, n=8) and incision in adults with neonatal incision (II, n=8).

### Early life pain (ELP) increases post injury evoked S1-mPFC connectivity in adult rats

The increased pain related signal processing in ELP found in adult S1, was not observed in mPFC. Since alterations in pain processing in mPFC may depend upon connections with other areas of the cerebral cortex, we next examined the functional connectivity between the S1 and mPFC in ELP rats. To explore this, we used phase locking value (PLV), a statistical method used to investigate task-induced changes in long range synchronization of neural activity (Lachaux et al., 1999) which provides an index of phase synchrony between two signals over a specific time period.

On the day of injury (D0), 2-3 hours after the incision, a significant increase in S1-mPFC PLV in response to eVF stimulation occurred in both ELP and non ELP rats following hindpaw skin incision (NI and II). There was no significant difference between the two injured groups (Figure6 A-C). This increase in phase locking was restricted to the S1 theta band and was not observed in other frequency bands (Delta: F_(2, 28)_ = 0.16, P=0.85 ; Alpha: F_(2, 28)_ = 1.75, P=0.19; Beta: F_(2, 28)_ = 0.96, P=0.39; Gamma: F_(2, 28)_ = 2.41, P=0.10). Importantly, a clear difference emerges upon inspection of the time course of this effect post injury, which reveals that the increased theta phase locking is maintained until 4 days post-injury in ELP (II) rats, compared to non-ELP (NI) groups (Figure 6D). We further examined correlation coefficients with pain behaviour to determine whether the increased S1-mPFC PLV in the theta band is associated with pain hypersensitivity. A significant inverse correlation between S1-mPFC PLV and PWT is seen in both in NI and II rats, but not in uninjured Con rats (Figure 6F).

**Figure 6:**
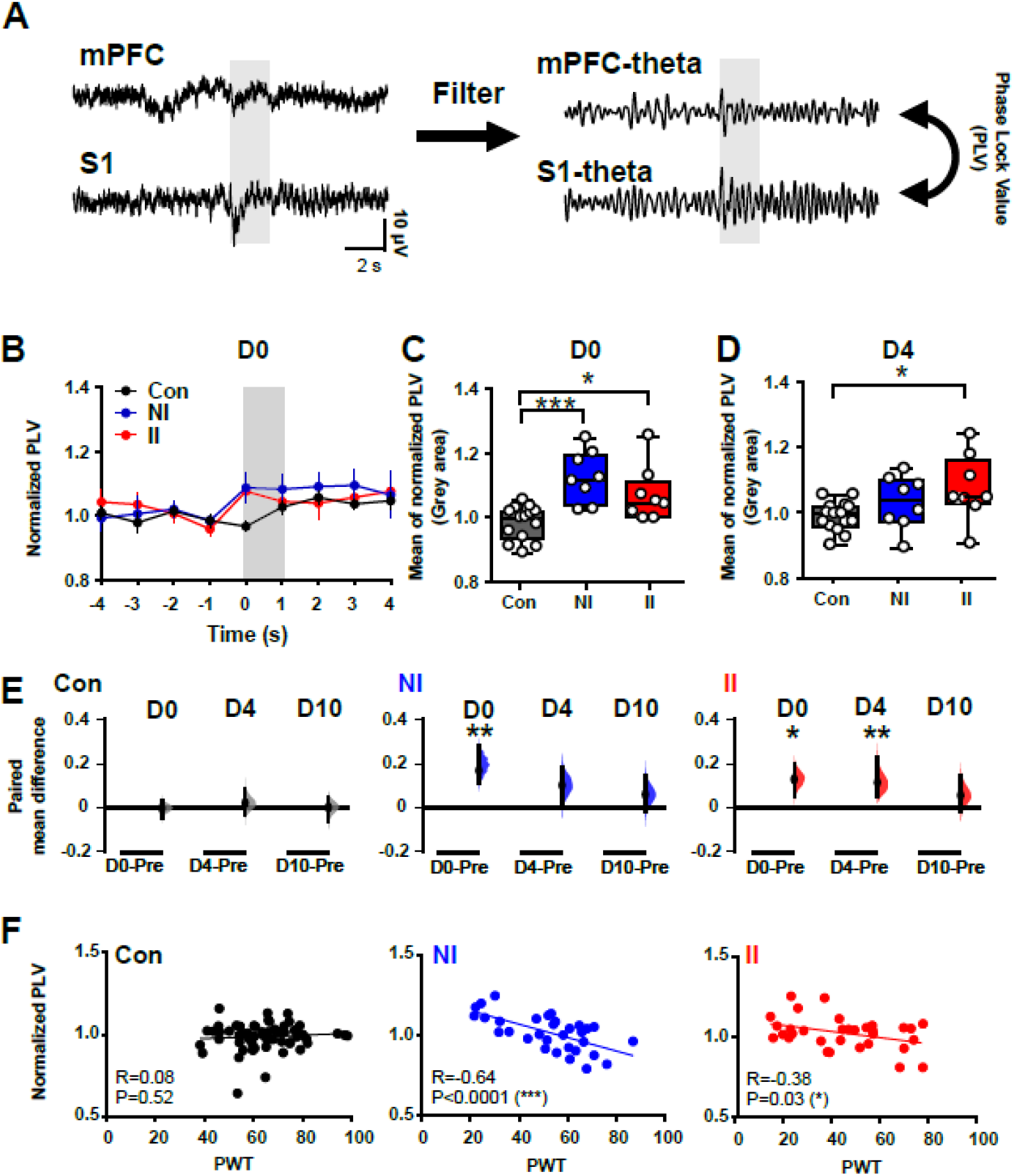
Early life pain (ELP) prolongs evoked beta phase coupling between S1 and mPFC post injury. Mechanical (eVF) stimulation of the hindpaw following injury in ELP adult rats (II, red), non-ELP rats (NI, blue) and controls (Con, black) while recording evoked potentials in the somatosensory cortex (S1). (A) The example of simultaneous traces recording of evoked LFPs from S1 and mPFC. (B) The S1-mPFC phase-locking value (PLV) at theta oscillations following injury (D0) were normalized to mean baseline (−4 to - 1 s) and present as normalized PLV (mean ± SEM). (C) The normalized PLV (D0/Pre) during evoked response as in Fig.3 (D) Evoked S1-mPFC PLV at theta frequencies at (D0), 4 days (D4) and 10 days (D10) after injury were compared to pre-injury (Pre) values. See Fig 3 for details.(E) Correlations between paw-withdraw threshold (PWT) and S1-mPFC phase lock theta oscillations in response to eVF. See Fig.3. for details *p < 0.05, **P<0.01, ***p < 0.001. Non-incised controls (Con, n=15), incision in adults without neonatal incision (NI, n=8) and incision in adults with neonatal incision (II, n=8).

## Discussion

The results presented here provide novel insights into the effects of ELP upon adult cortical pain networks. Using telemetric recording of local field potentials in the S1 and mPFC in awake adult mice we show that ELP results in significant changes in neural connectivity in the adult S1 and mPFC related to post-injury pain hypersensitivity.

We used a well-established model of ELP, skin incision on the plantar hindpaw, which when applied at a critical stage of development, is known to cause lasting changes in pain behaviour and increased post-injury pain hypersensitivity in adult life (Walker et al., 2009b; Beggs et al., 2012; Schwaller and Fitzgerald, 2014). The effect is driven and maintained by microglial activation in the dorsal horn of the spinal cord (Beggs et al., 2012; Moriarty et al., 2019) resulting in altered synaptic connectivity and reduced dynorphin inhibition (Li and Baccei, 2016, 2019; Brewer et al., 2020). Brainstem descending pain control is also altered in adults following early life incision (Walker et al., 2015) but the current data is the first to show changes in functional cortical pain networks following ELP. By recording simultaneous behavioural and cortical LFP responses to the same mechanical stimulus, we show that following ELP delta and gamma energy and delta/gamma modulation are increased in S1, together with increased phase-locking connectivity with mPFC, all directly correlated with behavioural pain hypersensitivity.

The result presented here provide new insight into the mechanisms whereby exposure to painful sensory experience in early life alters adult pain experience. The mPFC and S1 have key roles in cortical pain processing (Tan and Kuner, 2021); mPFC receives ascending nociceptive input, but also exerts important top-down regulation of sensory and affective processes of pain (Kummer et al., 2020), whereas S1 is the first level of pain perception and encodes nociceptive intensity and perceived pain intensity (Fields, 2012; Mancini et al., 2012). Pain is a complex phenomenon that depends on communication between different brain areas, which is served by neural oscillations and connectivity involving short-range and long-range communication processes (Baliki et al., 2011; Baliki and Apkarian, 2015; Kucyi and Davis, 2015; Ploner et al., 2017; Tan et al., 2021) and it is these oscillations that we have focussed on here.

The results show a significantly greater noxious-evoked gamma in S1 in injured rats with ELP. In humans, gamma-band oscillations in the primary somatosensory cortex correlate with subjective pain perception (Zhang et al., 2012; Heid et al., 2020) and in mice they are specifically strengthened, independently of any motor component, in the S1 cortex during nociception and are elevated during pain hypersensitivity (Tan et al., 2019). Nociceptive C fibre stimulation drives gamma activity in adult rat S1 (Chang et al., 2020b) and gamma oscillations generated by optogenetic activation of parvalbumin-expressing inhibitory interneurons in the S1 cortex enhance nociceptive sensitivity and induce aversive avoidance behaviour, while activating a network of prefrontal cortical and subcortical centres including descending serotonergic facilitatory pathways (Tan et al., 2021). Recent evidence suggests that gamma oscillations reflect strongly coupling of neural activity with fast spiking interneurons in the superficial layers of the S1 contralateral to the stimulated side (Yue et al., 2020). The increased energy of gamma oscillations, considered one of the most promising biomarkers of pain in the brain, is important evident for increased post injury pain perception in ELP animals.

Evoked activity in the delta frequency was also observed in the S1 of injured ELP rats. Event-related delta oscillations serve active sensory and cognitive functional roles across different sensory domains (Arnal and Giraud, 2012; Knyazev, 2012; Fardo et al., 2017) and play an crucial role in S1 sensory perception (Schroeder and Lakatos, 2009). Delta oscillations association with pain has been demonstrated elsewhere and may reflect coupling in thalamocortical loops (Sarnthein et al., 2006; Walton et al., 2010; Peng and Tang, 2016). The lack of delta frequency changes in mPFC supports the proposal that thalamo-S1 pathways are altered in ELP rats. Indeed, in human infants, ELP is associated with volume loss in the somatosensory thalamus accompanied by disruptions in thalamic metabolic growth and thalamocortical pathway maturation (Brummelte et al., 2012; Duerden et al., 2018).

Overall, these results indicate that the changes in delta and gamma activity in S1 are functionally linked to the behavioural hypersensitivity in injured rats with ELP. However, given the distinct intrinsic spatiotemporal properties of low- and high-frequency oscillations, we further examined the transient modulation of high-frequency amplitude (gamma) by low-frequency phase (delta) in relation to pain sensitivity and found enhanced evoked S1 delta-gamma modulation in injured rats with ELP. Because the high-frequency activity reflects local cortical processing, while low-frequency brain rhythms are dynamically entrained across distributed brain regions by both external sensory input and internal cognitive events, cross frequency modulation between low and high frequency is thought to contribute to information flow from large-scale brain networks to the fast, local cortical processing (Cardin et al., 2009; Canolty and Knight, 2010). Phase-amplitude cross-frequency coupling strength changes quickly in response to sensory, motor, and cognitive events (Schroeder and Lakatos, 2009) and abnormalities of cross frequency modulation may contribute to abnormal routing of information flow in chronic pain (Ploner et al., 2017). Our results suggest that such abnormal routing of information may occur in adults following ELP.

While the S1 reflects sensory discriminative aspects of pain, the prefrontal cortex is associated with the affective aspect of pain, providing top-down modulation of sensory and affective processes, including inhibition of both sensory and affective pain signals by descending projections to the various brain and spinal cord regions (Ji and Neugebauer, 2014; Bräscher et al., 2016; Kummer et al., 2020). Enhanced functional connectivity under procedural pain has been observed in several areas involved in pain perception: somatosensory cortices, anterior insula, anterior cingulate cortex and thalamus and mPFC (Bräscher et al., 2016; Galambos et al., 2019). Here we tested whether communication between S1 and mPFC was affected by ELP using synchronisation in the theta range as a measure of connectivity. Theta synchronization is proposed to be involved in large scale integration between long range multiple brain regions (von Stein and Sarnthein, 2000), especially in mPFC (Colgin, 2011; O’Neill et al., 2013; Esmaeili and Diamond, 2019), consistent with human data showing that prefrontal-sensorimotor connectivity is increased in tonic pain (Nickel et al., 2020). Our results show that adult skin injury does indeed produce a marked increase in evoked theta S1–mPFC connectivity, highly correlated to behavioural pain sensitivity in both ELP and control groups, but this increase is prolonged in ELP, lasting for 4 days compared to only one day in controls. Our data suggests that the connection between sensory and affective pain processing is enhanced in ELP rats which may underpin the wider social, emotional and cognitive life-long impact of ELP beyond increased pain perception (Ranger et al., 2018; de Kort et al., 2021; Ririe et al., 2021).

Our demonstration that ELP affects the cortical dynamics and connectivity underlying adult pain perception has important translational implications. Hospitalised infants exposed to ELP as a result of necessary clinical care, despite efforts to control that exposure (Laudiano-Dray et al., 2020; Eccleston et al., 2021), display long term structural and functional brain changes (Ranger and Grunau, 2014; Walker, 2019). Understanding the effects of ELP upon developing cortical pain networks will increase our understanding of individual susceptibility to pain in adult life (Denk et al., 2014).

## Acknowledgments

Supported by grants from the Biotechnology and Biological Sciences Research Council (MF, PC) (BB/R00823X/1) and the Medical Research Council (LF, MF) (MR/L019248/1).

## Author contributions

MF and PC conceived the study, PC collected the behavioural and electrophysiology data; PC and LF analysed the data; all authors interpreted the data and contributed to the manuscript.

## References

Apkarian AV, Bushnell MC, Treede R-D, Zubieta J-K (2005) Human brain mechanisms of pain perception and regulation in health and disease. European Journal of Pain 9:463–463.

Arnal LH, Giraud A-L (2012) Cortical oscillations and sensory predictions. Trends in Cognitive Sciences 16:390–398.

Baliki MN, Apkarian AV (2015) Nociception, Pain, Negative Moods, and Behavior Selection. Neuron 87:474–491.

Baliki MN, Baria AT, Apkarian AV (2011) The cortical rhythms of chronic back pain. J Neurosci 31:13981–13990.

Beggs S, Currie G, Salter MW, Fitzgerald M, Walker SM (2012) Priming of adult pain responses by neonatal pain experience: maintenance by central neuroimmune activity. Brain 135:404–417.

Bräscher A-K, Becker S, Hoeppli M-E, Schweinhardt P (2016) Different Brain Circuitries Mediating Controllable and Uncontrollable Pain. J Neurosci 36:5013.

Brennan TJ, Vandermeulen EP, Gebhart GF (1996) Characterization of a rat model of incisional pain. Pain 64: 493–502.

Brewer CL, Li J, O’Conor K, Serafin EK, Baccei ML (2020) Neonatal Injury Evokes Persistent Deficits in Dynorphin Inhibitory Circuits within the Adult Mouse Superficial Dorsal Horn. J Neurosci 40:3882–3895.

Brummelte S, Grunau RE, Chau V, Poskitt KJ, Brant R, Vinall J, Gover A, Synnes AR, Miller SP (2012) Procedural pain and brain development in premature newborns. Annals of Neurology 71:385–396.

Bruns A (2004) Fourier-, Hilbert-and wavelet-based signal analysis: are they really different approaches? Journal of Neuroscience Methods 137:321–332.

Buzsaki G (2004) Neuronal Oscillations in Cortical Networks. Science 304:1926–1929.

Canolty RT, Knight RT (2010) The functional role of cross-frequency coupling. Trends Cogn Sci 14:506–515.

Cao W, Lin S, Xia Q, Du Y, Yang Q, Zhang M, Lu Y, Xu J, Duan S, Xia J, Feng G, Xu J, Luo J (2018) Gamma Oscillation Dysfunction in mPFC Leads to Social Deficits in Neuroligin 3 R451C Knockin Mice. Neuron 97:1253–1260.e7.

Cardin JA, Carlén M, Meletis K, Knoblich U, Zhang F, Deisseroth K, Tsai L-H, Moore CI (2009) Driving fast-spiking cells induces gamma rhythm and controls sensory responses. Nature 459:663–667.

Chang P, Bush D, Schorge S, Good M, Canonica T, Shing N, Noy S, Wiseman FK, Burgess N, Tybulewicz VLJ, Walker MC, Fisher EMC (2020a) Altered Hippocampal-Prefrontal Neural Dynamics in Mouse Models of Down Syndrome. Cell Reports 30:1152–1163.e4.

Chang P, Fabrizi L, Fitzgerald M (2020b) Distinct Age-Dependent C Fiber-Driven Oscillatory Activity in the Rat Somatosensory Cortex. eNeuro 7:ENEURO.0036-20.2020.

Chang P, Fabrizi L, Olhede S, Fitzgerald M (2016) The Development of Nociceptive Network Activity in the Somatosensory Cortex of Freely Moving Rat Pups. Cerebral Cortex 26:4513–4523.

Chang P, Hashemi KS, Walker MC (2011) A novel telemetry system for recording EEG in small animals. Journal of Neuroscience Methods 201:106–115.

Colgin LL (2011) Oscillations and hippocampal–prefrontal synchrony. Current Opinion in Neurobiology 21:467–474.

de Kort AR, Joosten EA, Patijn J, Tibboel D, van den Hoogen NJ (2021) Neonatal procedural pain affects state, but not trait anxiety behavior in adult rats. Dev Psychobiol 63:e22210.

Denk F, McMahon SB, Tracey I (2014) Pain vulnerability: a neurobiological perspective. Nat Neurosci 17:192–200.

Duerden EG, Albanese M-C (2013) Localization of pain-related brain activation: A meta-analysis of neuroimaging data. Human Brain Mapping 34:109–149.

Duerden EG, Grunau RE, Guo T, Foong J, Pearson A, Au-Young S, Lavoie R, Chakravarty MM, Chau V, Synnes A, Miller SP (2018) Early Procedural Pain Is Associated with Regionally-Specific Alterations in Thalamic Development in Preterm Neonates. J Neurosci 38:878–886.

Eccleston C et al. (2021) Delivering transformative action in paediatric pain: a Lancet Child & Adolescent Health Commission. Lancet Child Adolesc Health 5:47–87.

Esmaeili V, Diamond ME (2019) Neuronal Correlates of Tactile Working Memory in Prefrontal and Vibrissal Somatosensory Cortex. Cell Reports 27:3167–3181.e5.

Euston DR, Gruber AJ, McNaughton BL (2012) The role of medial prefrontal cortex in memory and decision making. Neuron 76:1057–1070.

Fardo F, Vinding MC, Allen M, Jensen TS, Finnerup NB (2017) Delta and gamma oscillations in operculo-insular cortex underlie innocuous cold thermosensation. J Neurophysiol 117:1959–1968.

Ferrier J, Bayet-Robert M, Dalmann R, El Guerrab A, Aissouni Y, Graveron-Demilly D, Chalus M, Pinguet J, Eschalier A, Richard D, Daulhac L, Marchand F, Balayssac D (2015) Cholinergic Neurotransmission in the Posterior Insular Cortex Is Altered in Preclinical Models of Neuropathic Pain: Key Role of Muscarinic M2 Receptors in Donepezil-Induced Antinociception. J Neurosci 35:16418.

Ferrier J, Marchand F, Balayssac D (2016) Assessment of Mechanical Allodynia in Rats Using the Electronic Von Frey Test. Bio-protocol 6:e1933.

Fields HL (2012) Pain and the primary somatosensory cortex. Pain 153:742–743.

Florin E, Baillet S (2015) The brain’s resting-state activity is shaped by synchronized cross-frequency coupling of neural oscillations. Neuroimage 111:26–35.

Galambos A, Szabó E, Nagy Z, Édes AE, Kocsel N, Juhász G, Kökönyei G (2019) A systematic review of structural and functional MRI studies on pain catastrophizing. J Pain Res 12:1155–1178.

Heid C, Mouraux A, Treede R-D, Schuh-Hofer S, Rupp A, Baumgärtner U (2020) Early gamma-oscillations as correlate of localized nociceptive processing in primary sensorimotor cortex. Journal of Neurophysiology 123:1711–1726.

Ho J, Tumkaya T, Aryal S, Choi H, Claridge-Chang A (2019) Moving beyond P values: data analysis with estimation graphics. Nature Methods 16:565–566.

Huang W-C, Zucca A, Levy J, Page DT (2020) Social Behavior Is Modulated by Valence-Encoding mPFC-Amygdala Sub-circuitry. Cell Reports 32:107899.

Ji G, Neugebauer V (2014) CB1 augments mGluR5 function in medial prefrontal cortical neurons to inhibit amygdala hyperactivity in an arthritis pain model. Eur J Neurosci 39:455–466.

Jones GT, Power C, Macfarlane GJ (2009) Adverse events in childhood and chronic widespread pain in adult life: Results from the 1958 British Birth Cohort Study. Pain 143:92–96.

Jones L, Verriotis M, Cooper RJ, Laudiano-Dray MP, Rupawala M, Meek J, Fabrizi L, Fitzgerald M (2021) Widespread nociceptive maps in the human neonatal somatosensory cortex. :2021.07.29.454164 https://www.biorxiv.org/content/10.1101/2021.07.29.454164v1

Knyazev GG (2012) EEG delta oscillations as a correlate of basic homeostatic and motivational processes. Neuroscience & Biobehavioral Reviews 36:677–695.

Kramer MA, Tort ABL, Kopell NJ (2008) Sharp edge artifacts and spurious coupling in EEG frequency comodulation measures. Journal of Neuroscience Methods 170:352–357.

Kucyi A, Davis KD (2015) The dynamic pain connectome. Trends Neurosci 38:86–95.

Kummer KK, Mitrić M, Kalpachidou T, Kress M (2020) The Medial Prefrontal Cortex as a Central Hub for Mental Comorbidities Associated with Chronic Pain. Int J Mol Sci 21:3440.

Lachaux J-P, Rodriguez E, Martinerie J, Varela FJ (1999) Measuring phase synchrony in brain signals. Human Brain Mapping 8:194–208.

Laudiano-Dray MP, Pillai Riddell R, Jones L, Iyer R, Whitehead K, Fitzgerald M, Fabrizi L, Meek J (2020) Quantification of neonatal procedural pain severity: a platform for estimating total pain burden in individual infants. Pain 161:1270–1277.

Le Van Quyen M, Foucher J, Lachaux J-P, Rodriguez E, Lutz A, Martinerie J, Varela FJ (2001) Comparison of Hilbert transform and wavelet methods for the analysis of neuronal synchrony. Journal of Neuroscience Methods 111:83–98.

Li J, Baccei ML (2016) Neonatal Tissue Damage Promotes Spike Timing-Dependent Synaptic Long-Term Potentiation in Adult Spinal Projection Neurons. J Neurosci 36:5405–5416.

Li J, Baccei ML (2019) Neonatal Injury Alters Sensory Input and Synaptic Plasticity in GABAergic Interneurons of the Adult Mouse Dorsal Horn. J Neurosci 39:7815.

Mancini F, Haggard P, Iannetti GD, Longo MR, Sereno MI (2012) Fine-grained nociceptive maps in primary somatosensory cortex. J Neurosci 32:17155–17162.

Melchior M, Kuhn P, Poisbeau P (2021) The burden of early life stress on the nociceptive system development and pain responses. Eur J Neurosci.

Moriarty O, Tu Y, Sengar AS, Salter MW, Beggs S, Walker SM (2019) Priming of adult incision response by early life injury: neonatal microglial inhibition has persistent but sexually dimorphic effects in adult rats. J Neurosci:1786–18.

Nickel MM, Ta Dinh S, May ES, Tiemann L, Hohn VD, Gross J, Ploner M (2020) Neural oscillations and connectivity characterizing the state of tonic experimental pain in humans. Human Brain Mapping 41:17–29.

O’Neill P-K, Gordon JA, Sigurdsson T (2013) Theta Oscillations in the Medial Prefrontal Cortex Are Modulated by Spatial Working Memory and Synchronize with the Hippocampus through Its Ventral Subregion. The Journal of Neuroscience 33:14211–14224.

Ong W-Y, Stohler CS, Herr DR (2019) Role of the Prefrontal Cortex in Pain Processing. Mol Neurobiol 56:1137–1166.

Paxinos G, Ashwell KWS, Tork I (2013) Atlas of the Developing Rat Nervous System. Academic Press.

Peng W, Tang D (2016) Pain Related Cortical Oscillations: Methodological Advances and Potential Applications. Frontiers in Computational Neuroscience 10:9.

Ploner M, Sorg C, Gross J (2017) Brain Rhythms of Pain. Trends Cogn Sci 21:100–110.

Ranger M, Grunau RE (2014) Early repetitive pain in preterm infants in relation to the developing brain. Pain Manag 4:57–67.

Ranger M, Tremblay S, Chau CMY, Holsti L, Grunau RE, Goldowitz D (2018) Adverse Behavioral Changes in Adult Mice Following Neonatal Repeated Exposure to Pain and Sucrose. Front Psychol 9:2394.

Ririe DG, Eisenach JC, Martin TJ (2021) A Painful Beginning: Early Life Surgery Produces Long-Term Behavioral Disruption in the Rat. Front Behav Neurosci 15:630889.

Samiee S, Baillet S (2017) Time-resolved phase-amplitude coupling in neural oscillations. NeuroImage 159:270–279.

Sarnthein J, Stern J, Aufenberg C, Rousson V, Jeanmonod D (2006) Increased EEG power and slowed dominant frequency in patients with neurogenic pain. Brain 129:55–64.

Schroeder CE, Lakatos P (2009) Low-frequency neuronal oscillations as instruments of sensory selection. Trends Neurosci 32:9–18.

Schwaller F, Fitzgerald M (2014) The consequences of pain in early life: injury-induced plasticity in developing pain pathways. Eur J Neurosci 39:344–352.

Tadel F, Baillet S, Mosher JC, Pantazis D, Leahy RM (2011) Brainstorm: A User-Friendly Application for MEG/EEG Analysis Oostenveld R, ed. Computational Intelligence and Neuroscience 2011:879716.

Tan LL, Kuner R (2021) Neocortical circuits in pain and pain relief. Nat Rev Neurosci 22:458–471.

Tan LL, Oswald MJ, Heinl C, Retana Romero OA, Kaushalya SK, Monyer H, Kuner R (2019) Gamma oscillations in somatosensory cortex recruit prefrontal and descending serotonergic pathways in aversion and nociception. Nature Communications 10:983.

Tan LL, Oswald MJ, Kuner R (2021) Neurobiology of brain oscillations in acute and chronic pain. Trends Neurosci 44:629–642.

Torsney C, Fitzgerald M (2003) Spinal dorsal horn cell receptive field size is increased in adult rats following neonatal hindpaw skin injury. J Physiol (Lond) 550:255–261.

van den Hoogen NJ, Patijn J, Tibboel D, Joosten BA, Fitzgerald M, Kwok CHT (2018) Repeated touch and needle-prick stimulation in the neonatal period increases the baseline mechanical sensitivity and postinjury hypersensitivity of adult spinal sensory neurons. Pain 159:1166–1175.

von Stein A, Sarnthein J (2000) Different frequencies for different scales of cortical integration: from local gamma to long range alpha/theta synchronization. International Journal of Psychophysiology 38:301–313.

Walker SM (2019) Long-term effects of neonatal pain. Semin Fetal Neonatal Med 24:101005.

Walker SM, Fitzgerald M, Hathway GJ (2015) Surgical injury in the neonatal rat alters the adult pattern of descending modulation from the rostroventral medulla. Anesthesiology 122:1391–1400.

Walker SM, Franck LS, Fitzgerald M, Myles J, Stocks J, Marlow N (2009a) Long-term impact of neonatal intensive care and surgery on somatosensory perception in children born extremely preterm. Pain 141:79–87.

Walker SM, O’Reilly H, Beckmann J, Marlow N, EPICure@19 Study Group (2018) Conditioned pain modulation identifies altered sensitivity in extremely preterm young adult males and females. Br J Anaesth 121:636–646.

Walker SM, Tochiki KK, Fitzgerald M (2009b) Hindpaw incision in early life increases the hyperalgesic response to repeat surgical injury: Critical period and dependence on initial afferent activity. Pain 147: 99–106.

Walton KD, Dubois M, Llinás RR (2010) Abnormal thalamocortical activity in patients with Complex Regional Pain Syndrome (CRPS) Type I. Pain 150: 41–51.

Yue L, Iannetti GD, Hu L (2020) The Neural Origin of Nociceptive-Induced Gamma-Band Oscillations. J Neurosci 40:3478–3490.

Zhang Z, Gadotti VM, Chen L, Souza IA, Stemkowski PL, Zamponi GW (2015) Role of Prelimbic GABAergic Circuits in Sensory and Emotional Aspects of Neuropathic Pain. Cell Reports 12:752–759.

Zhang ZG, Hu L, Hung YS, Mouraux A, Iannetti GD (2012) Gamma-band oscillations in the primary somatosensory cortex--a direct and obligatory correlate of subjective pain intensity. J Neurosci 32:7429–7438.

